# A deep learning and graph-based approach to characterise the immunological landscape and spatial architecture of colon cancer tissue

**DOI:** 10.1101/2022.07.06.498984

**Authors:** Mario Parreno-Centeno, Guidantonio Malagoli Tagliazucchi, Eloise Withnell, Shi Pan, Maria Secrier

## Abstract

Tumour immunity is key for the prognosis and treatment of colon adenocarcinoma, but its characterisation remains cumbersome and expensive, requiring sequencing or other complex assays. Detecting tumour-infiltrating lymphocytes in haematoxylin and eosin (H&E) slides of cancer tissue would provide a cost-effective alternative to support clinicians in treatment decisions, but inter- and intra-observer variability can arise even amongst experienced pathologists. Furthermore, the compounded effect of other cells in the tumour microenvironment is challenging to quantify but could yield useful additional biomarkers. We combined RNA sequencing, digital pathology and deep learning through the InceptionV3 architecture to develop a fully automated computer vision model that detects prognostic tumour immunity levels in H&E slides of colon adenocarcinoma with an area under the curve (AUC) of 82%. Amongst tumour infiltrating T cell subsets, we demonstrate that CD8+ effector memory T cell patterns are most recognisable algorithmically with an average AUC of 83%. We subsequently applied nuclear segmentation and classification via HoVer-Net to derive complex cell-cell interaction graphs, which we queried efficiently through a bespoke Neo4J graph database. This uncovered stromal barriers and lymphocyte triplets that could act as structural hallmarks of low immunity tumours with poor prognosis. Our integrated deep learning and graph-based workflow provides evidence for the feasibility of automated detection of complex immune cytotoxicity patterns within H&E-stained colon cancer slides, which could inform new cellular biomarkers and support treatment management of this disease in the future.

## INTRODUCTION

Tumour immunity is a critical determinant of clinical outcome in colon adenocarcinoma^1,2^, yet its characterisation is challenging due to cellular heterogeneity and the elaborate techniques required to study it comprehensively^3^. Earlier efforts introduced the prognostic value of the CD8+ to CD4+ T cell infiltration ratio within the tumour^4,5^. Galon et al^6^ built on this to develop an ‘Immunoscore’ that predicts the risk of recurrence and chemotherapy response in colorectal cancer^7,8^. This score was derived based on immunohistochemistry (IHC), a fairly elaborate procedure to label antigens expressed by specific cells, which requires well defined markers and is not incorporated routinely in the clinical workflow. Instead, H&E staining of nucleus, cytoplasm and extracellular matrix offers a cheaper, streamlined alternative in oncology. Experienced pathologists can identify tumour infiltrating lymphocytes and other cells in H&E-stained tissue, but inter-and intra-observer variability can arise when assessing a sample^9^. Therefore, automating the detection of such cells could prove beneficial^10,11^. Indeed, deep learning algorithms have recently been employed for this purpose in multiple cancers^12-14^, with links to patient outcome demonstrated in colorectal cancer^2,15^.

However, these models often focus on individual cell types and do not consider the broader spatial organisation of the tissue, which can impact tumour growth and dissemination^16^. For instance, the stroma can promote tumour progression by limiting access of therapeutic agents or immune cells to tumours through fibrosis^17^. Such factors are currently challenging to assess systematically because of costs and a lack of standardised methodology. Several studies have shown that spatial organisation features such as the stromal content/ratios, TIL morphology and even immune hot/cold phenotypes can be extracted from H&E patches and linked to prognosis^12,18-23^. AbdulJabbar et al^24^ and Bilal et al^25^ have also integrated deep learning methods on digital histology and omics data more extensively, gaining insights into how cellular organisation impacts cancer evolution and clinical outcomes. However, these studies often require expert annotation and still lack insight into more complex interactions between different cell types.

As an alternative to expert annotation, developments of computational tools for immune deconvolution from RNA sequencing have revolutionised our ability to capture an extensive array of cell types and states in bulk tissue^26-28^. Cancer transcriptomics datasets are widely available and have been successfully integrated with H&E images via deep learning in recent studies, demonstrating that patterns of angiogenesis, hypoxia and even T and B cell immunity are detectable in the tissue^29^. Integrating such approaches to characterise tumour immunity as a whole within the colon cancer tissue would be highly informative for prognosis and treatment, but has not yet been achieved.

This study tests the extent to which tumour immunity and its spatial organisation can be quantified within colon adenocarcinoma tissue based on transcriptomic signatures alone. We propose a fully automated image-processing pipeline to predict the immune activity in a tumour from H&E images based on matched bulk RNA-seq data. We also introduce a novel framework for surveying detailed cellular interaction landscapes within digital pathology slides by combining nuclear segmentation, classification and graph assembly, with efficient queries handled via a bespoke Neo4J graph database. Finally, we apply this framework to interrogate the complex tissue organisation in colon adenocarcinoma and identify cellular structures linked with immune response, which we then validate using spatial transcriptomics.

## RESULTS

### Transcriptome-derived tumour immunity informs prognosis, molecular characteristics and treatment efficacy in colon adenocarcinoma

To explore the immune landscape and tissue organisation of colon adenocarcinoma, we employed RNA sequencing (RNA-seq) and matched digital pathology data collected from tumour samples of 456 patients available from TCGA. We calculated a tumour immunity score per sample as the average abundance of tumour infiltrating cells from the microenvironment using ConsensusTME^28^ (Supplementary Figure 1a). This score accounted for the compounded effects of expression signals coming from all detectable non-tumour cells, reflecting a spectrum of low to high immunity (Supplementary Figure 1b). The score correlated well with the mean CD8+/CD4+ T cell subtype scores estimated from transcriptomic data using an alternative immune deconvolution method, xCell (R = 0.46, p<2e-16).

After quantifying the overall tumour immunity in each sample, we assessed the prognostic value of the respective immune scores. To maximise the clinical relevance of this analysis, we sought to identify a tumour immunity score threshold that would reflect the strongest relation with patient outcome. We found that a cut-off point of 0·39 highlighted two immunity groups (High Immunity, HI, and Low Immunity, LI) with highly distinct overall survival (p=0·002, Figure 1b-c) and disease-free intervals (p=0·005, Supplementary Figure 1c). As expected, the patients with high levels of tumour immunity showed significantly better outcomes. When checking against the pathology annotations of the slides, the differences between the HI and LI groups appeared to be driven by the presence of lymphocytes and neutrophils, which were elevated in the HI group (Figure 1d). Furthermore, the HI tumours also had elevated markers of intratumoural natural killer (NK) cell activity, including expression of NK cell receptors and corresponding tumour cell ligands, as well as secreted cytokines (Supplementary Figure 1d), which have been associated with immunotherapy response^30^.

**Figure 1.**
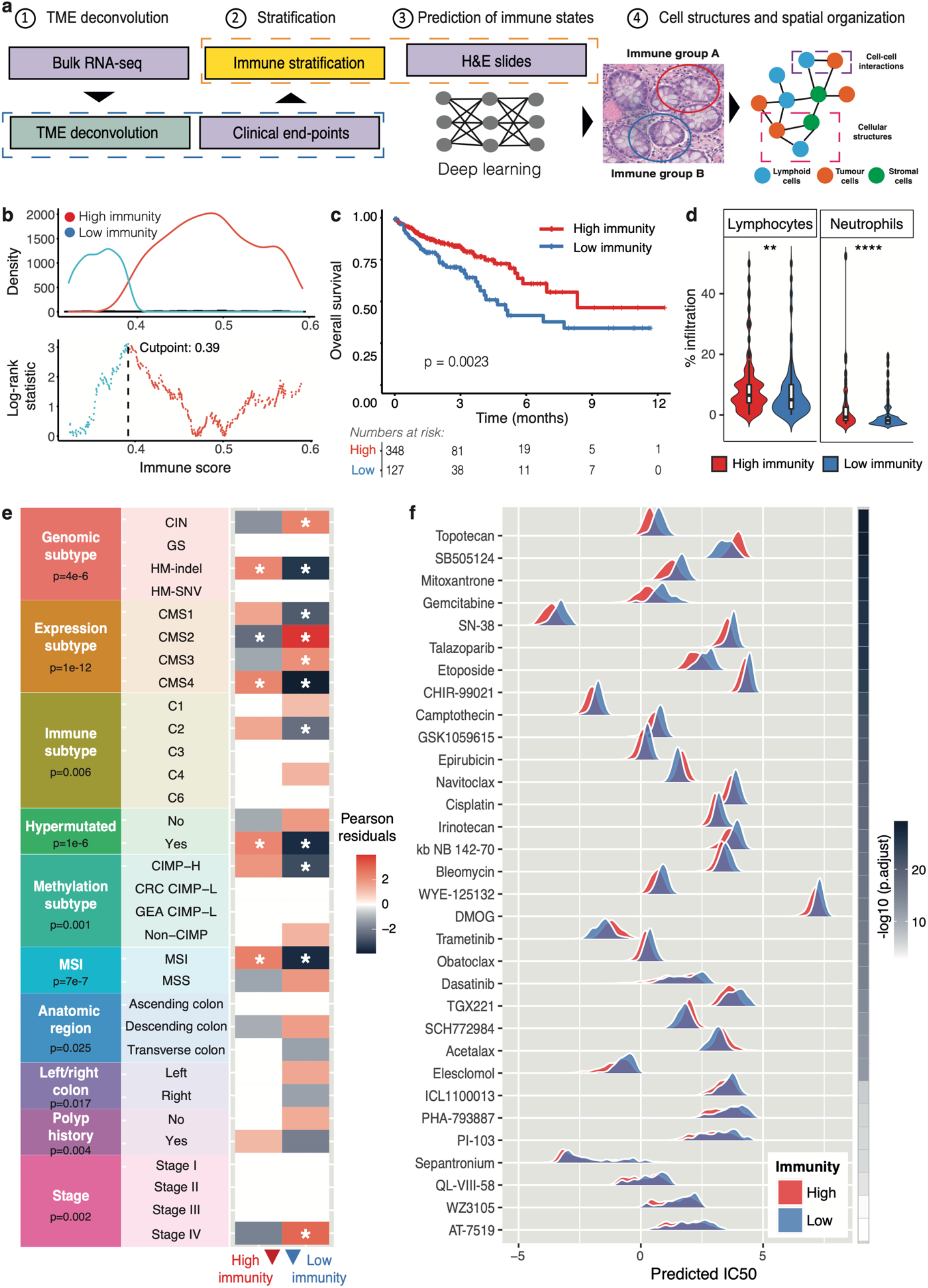
Study workflow and immunity-based stratification of colon adenocarcinoma. **(a)** Workflow of the study. A prognostic classifier of colon adenocarcinoma was defined based on RNA-seq inferred immune activity in the tumours. An H&E based deep learning classifier was then on these labels, and differences in cellular compositions and interactions between the two groups were subsequently described. **(b)** Optimisation of tumour immunity threshold to maximise survival differences. The dotted line highlights the optimal cut-point of 0.39. **(c)** The high and low immunity groups defined using the cut-off in (b) show significantly different overall survival in the TCGA-COAD cohort. (d) The high immunity group presents higher fractions of lymphocytes and neutrophils, as scored by pathologists. (e) Differences in colon adenocarcinoma molecular subgroup and clinical characteristics between high and low immunity tumours, inferred from conditional independence tests. Only significantly associated characteristics are shown. The stars mark Pearson residuals greater than 2 or less than -2, indicating the strongest correlations. **(f)** Predicted drug sensitivity of TCGA-COAD tumours to a variety of anti-cancer compounds, compared between high and low immunity groups. Only compounds showing significant differences in drug sensitivity are shown (ranked by the magnitude of the difference).

Colon cancer has been systematically characterised from a genomic point of view, with mutation, expression and methylation features shown to associate with disease progression and therapy responses^31,32^. To link these findings with our classification, we further characterised the 456 samples in the HI/LI groups using molecular and tumour architecture phenotypes already described for colon adenocarcinomas^31^ (Figure 1e). Genomically, we found that low immunity tumours tended to fall within the chromosomally unstable group of gastrointestinal cancers (CIN). Indeed, tumours with higher levels of CIN/aneuploidy have been previously linked with immune evasion and poorer outcomes to immunotherapies^33^. In contrast, high immunity tumours were associated with the gastrointestinal hypermutated indel (HM-indel) subtype and presented frequent microsatellite instability (MSI), in line with numerous studies linking hypermutation with increased neoantigen presentation and activation of the host antitumor immune response^34,35^.

The consensus expression-based classification defined by Guinney et al^32^ revealed that the high immunity tumours tended to be more mesenchymal (CMS4), a subtype characterised by increased stromal infiltration and angiogenesis. In contrast, lower immunity was linked with epithelial, metabolically deregulated phenotypes (CMS2/3), and lacked the characteristic pattern of CpG island hypermethylation (non-CIMP). Anatomically, low immunity was more frequently characteristic of tumours arising in the descending colon and on the left side, and prevailed in late-stage cancers, where immune recognition is more likely impaired. Tumours with higher immunity tended to present fewer hyperplastic polyps.

Finally, the two defined immunity groups in colon adenocarcinomas were predicted to display significantly different susceptibility to multiple anti-cancer compounds (Figure 1f). The largest differences were observed for chemotherapy drugs such as topotecan, mitoxantrone and gemcitabine. Higher immune activity was linked with lower IC50 values, suggesting greater efficacy in this setting and confirming previous links between immunity and enhanced chemotherapeutic responses^8^.

### A prognostic classifier of tumour immunity in colon cancer from H&E-stained images

Having demonstrated the clinical relevance of our immune classification of colon adenocarcinomas, we then developed a digital pathology classifier of this phenotype. We trained a deep learning model based on the InceptionV3 architecture to classify high and low tumour immunity (as inferred from matched RNA-seq data) in H&E-stained images of cancer tissue collected from colon adenocarcinoma patients (Figure 2a). We achieved a median accuracy of 82% AUC probability based on tile average and percentage count in the testing dataset (Fig 2b-c). This approach also allowed us to obtain a spatially-resolved overview of tumour immunity levels within entire tissue slides through tile-level estimates (Fig 2d). Indeed, tissue slides from highly immunogenic tumours were predominantly composed of tiles with strong HI signals, whereas low immunity tumours presented more diverse predictions of LI and intermediate levels of immunity across tiles (Fig 2e).

**Figure 2.**
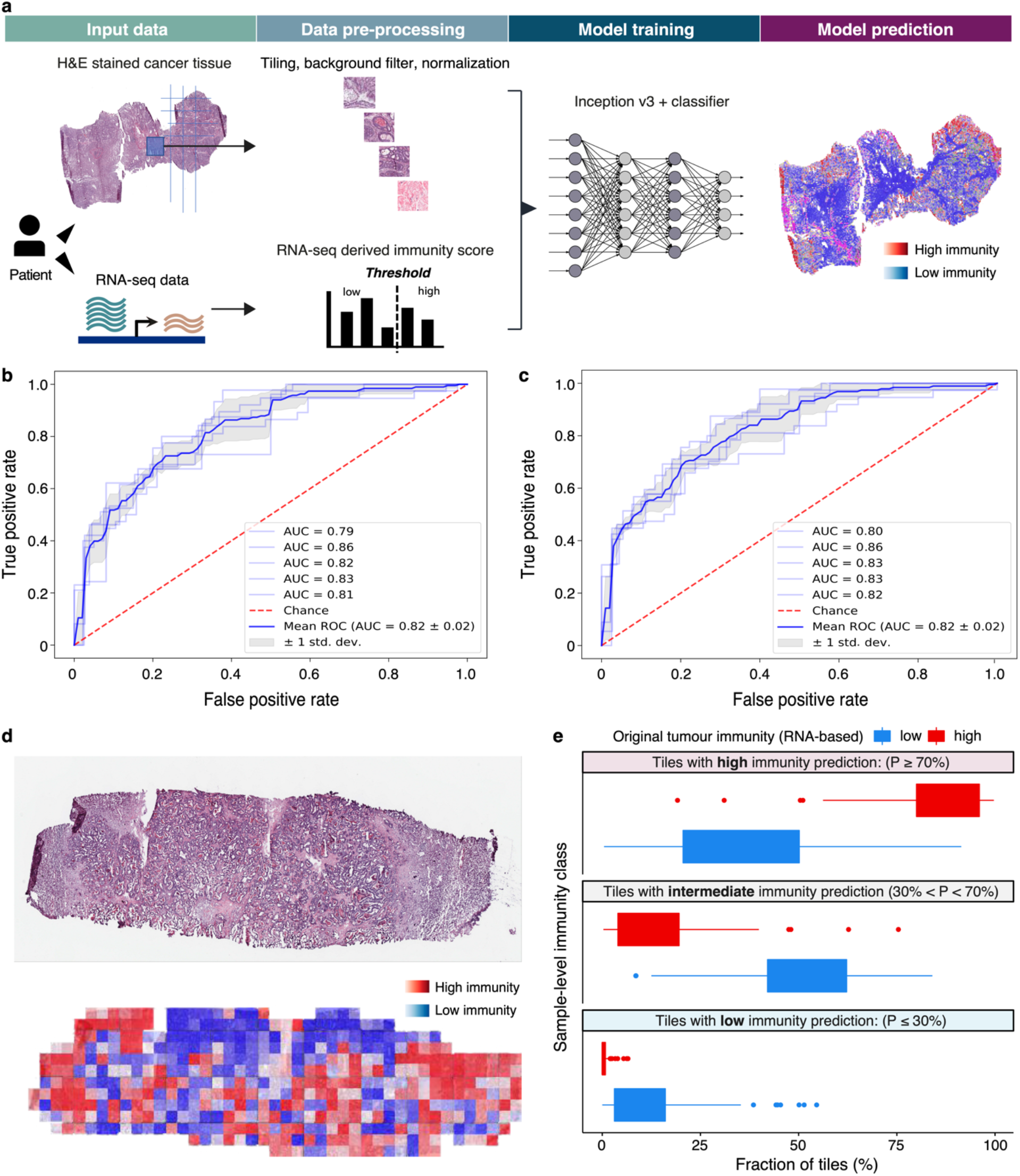
Deep learning classifier of tumour immunity. (a) Pipeline for the identification of RNA-seq based immune phenotypes in digital images of H&E stained cancer tissue. (b-c) A mean accuracy (AUC) of 82% is obtained when predicting the immune phenotype of colon cancer H&E slides with 5-fold cross validation by averaging the probability per tile (b) and counting the percentage of tiles (c), respectively. (d) Example of an H&E stained slide (top) and the corresponding immunity predictions of the model within the same slide (bottom). The colour gradient from blue to red reflects increasing probability of high immune content in each patch. (e) Immunity class deep learning predictions in each tile, compared between samples with high and low overall immunity (as inferred from RNA-seq data). Tile-level predictions have been classed as high, intermediate or low immunity based on the probability of belonging to the HI/LI group as indicated.

While the HI group of tumours is expected to have enhanced immune reactivity, the cytotoxic effect is triggered by populations of CD8+ and CD4+ immune cells that are transformed from a naïve to an effector state upon antigen recognition^36^. These CD8+/CD4+ subsets emerge at different points of cancer development and could create distinct patterns within the tumour tissue. The ability of deep learning algorithms to distinguish these immune subsets in H&E slides has not been tested in this system. To understand whether patterns of active rather than uneducated immunity might be preferentially captured, we trained deep learning models to identify distinct subgroups of CD4+ and CD8+ T cells as characterised in Aran et al^27^, i.e., CD4/CD8+ naïve T cells, central memory cells and effector memory cells in the TCGA COAD cohort. Our models could detect expression signals of CD4+ central memory T cells and CD8+ effector memory T cells with high accuracies of 82% and 83% AUC, respectively (Figure 3). The rest of the cells were identified with lower accuracy, between 64% and 70% AUC. Overall, this suggests that the cells most relevant to triggering an effective immune response are also the ones that leave the most recognisable traces within the colon cancer tissue and could be specifically tracked in a clinical setting to support immunotherapy treatment decisions.

**Figure 3.**
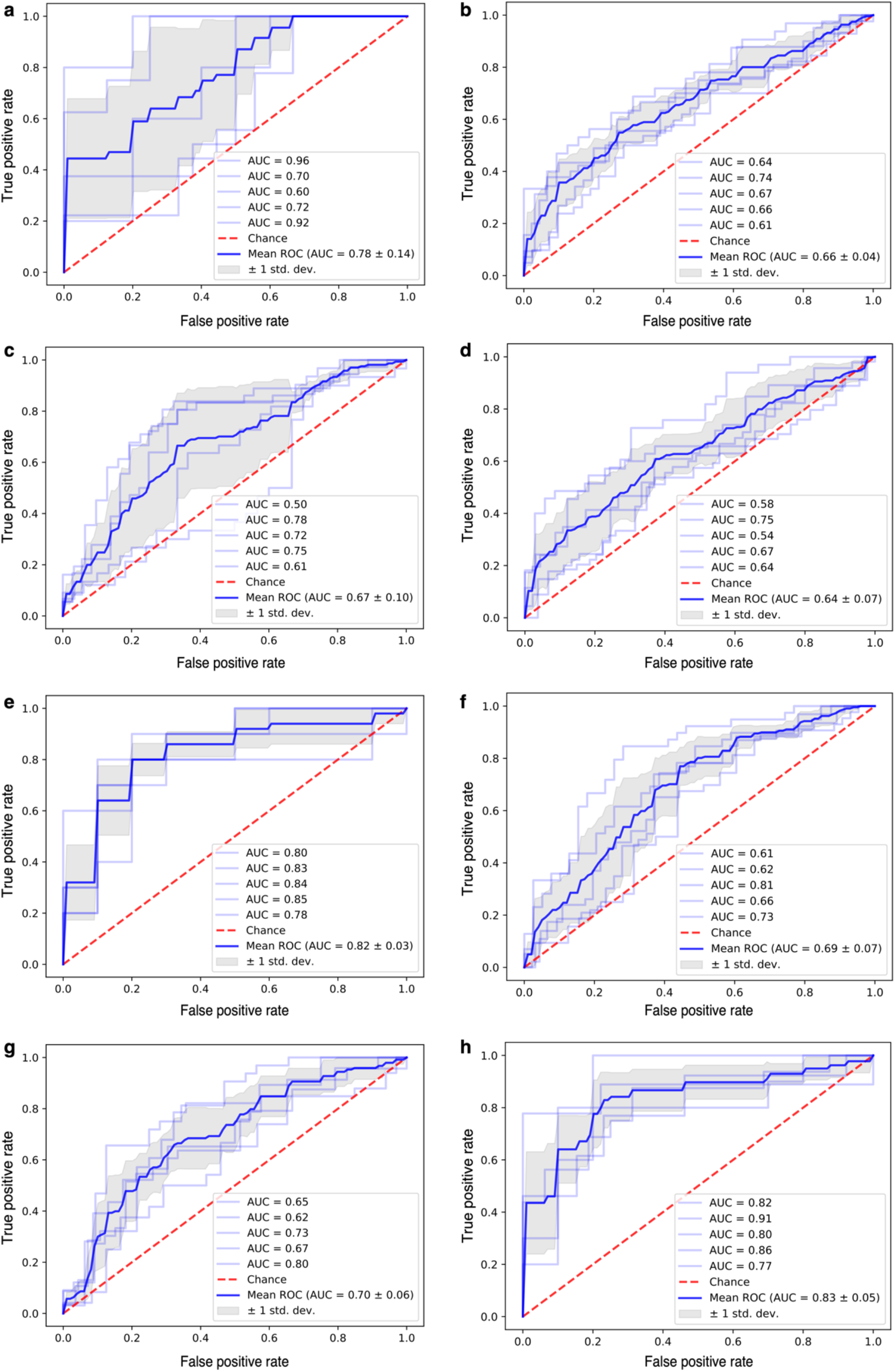
Prediction of the subgroups of CD4+ and CD8+ T cells in H&E images. Mean accuracies (AUC) are shown for predictions of each of the following T cell categories: (a) generic CD4+ T cells - 78%; (b) generic CD8+ T cells - 66%; (c) CD4+ naïve T cells - 67%; (d) CD8+ naïve T cells – 64%; (e) CD4+ central memory T cells - 82%; (f) CD8+ central memory T cells - 69%; (g) CD4+ effector memory T cells - 70%; (h) CD8+ effector memory T cells - 83%.

### Cellular organisation of the immune response in colon adenocarcinoma

Next, we sought to gain further insight into the cellular organisation of colon adenocarcinoma tissue in a high versus low immunity setting using the H&E slide information. Using HoVer-Net^37^, we conducted nuclear segmentation and classification within the H&E tiles of the TCGA COAD dataset. This approach allowed us to label individual cells within the tissue as ‘tumour’, ‘lymphocytic’ or ‘stromal’ (Figure 4a). Based on cell proximity, we could then infer interactions between different cells and summarise them for every tissue patch in the form of graphs (Figure 4a). However, this analysis resulted in thousands to hundreds of thousands of interactions per sample, yielding complex graphs that are difficult to store, integrate and query. To resolve this, we developed a Neo4J graph database comprising 2,786,464 nodes and 3,628,377 edges, that would allow us to explore and interrogate such complex structures effectively. The basic relationship model in the database is depicted in Figure 4b. This database allows us to perform complex queries to identify biologically relevant structures, such as a stromal barrier separating lymphocytes from tumour cells (Figure 4c), or a lymphocytic attack on cancer cells (Figure 4d). These graph interactions can also be dynamically explored and expanded within the database (Supplementary Video 1).

**Figure 4.**
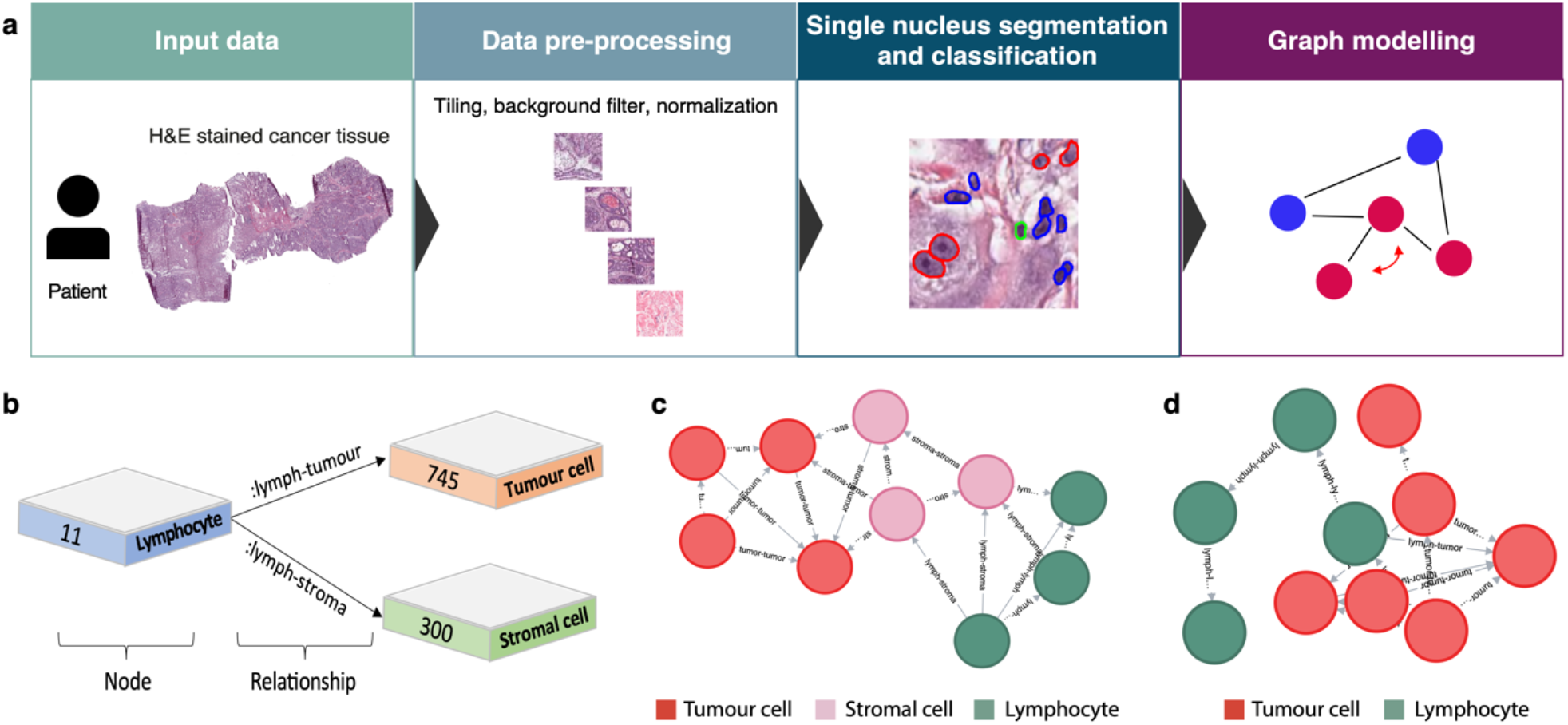
Graph analysis of the cellular organisation of the immune response in colon adenocarcinoma. (a) Pipeline of the graph analysis approach. After preprocessing the WSIs, cells are segmented and classified using HoVer-Net, following further graph modelling and storage. (b) The inferred cell-cell interactions within the tissue are stored in a Neo4j graph database for further analysis. The database structure is shown, with representative relationships between cell types illustrated. (c) Example of a stromal barrier graph structure, separating tumour cells from lymphocytes. (d) Example of a graph structure illustrating lymphocyte infiltration of the tumour.

We compared the cell population frequencies and interactions between the HI and LI groups. Lymphocytes and stromal cells were similarly abundant in the two groups, but appeared more frequently isolated in a high immunity context (Figure 5a-b). These could correspond to exhausted or naïve T cells being recruited to the site, which would be less likely to form interactions. As expected, the HI group also harboured more frequent direct interactions between lymphocytes and tumour cells, suggesting immune recognition typical of ‘immune hot’ phenotypes^38^ (Figure 5c,e). The number of tumour-stroma interactions was also increased (Figure 5d), but to a lower extent (Figure 5e). On the other hand, dense lymphocyte clusters in the form of triplets, as well as stromal barriers were increased in the low immunity tumours (Figure 5f-g). The lymphocyte triplet presence might suggest inactive clusters that are not recognising malignant cells. Concurrently, the stromal barriers may aid immune evasion, as has been previously described in pancreatic cancer^39^. These cellular structures reflect different cellular organisation and immune activity in an HI vs LI setting and could support treatment strategies in the clinic.

**Figure 5.**
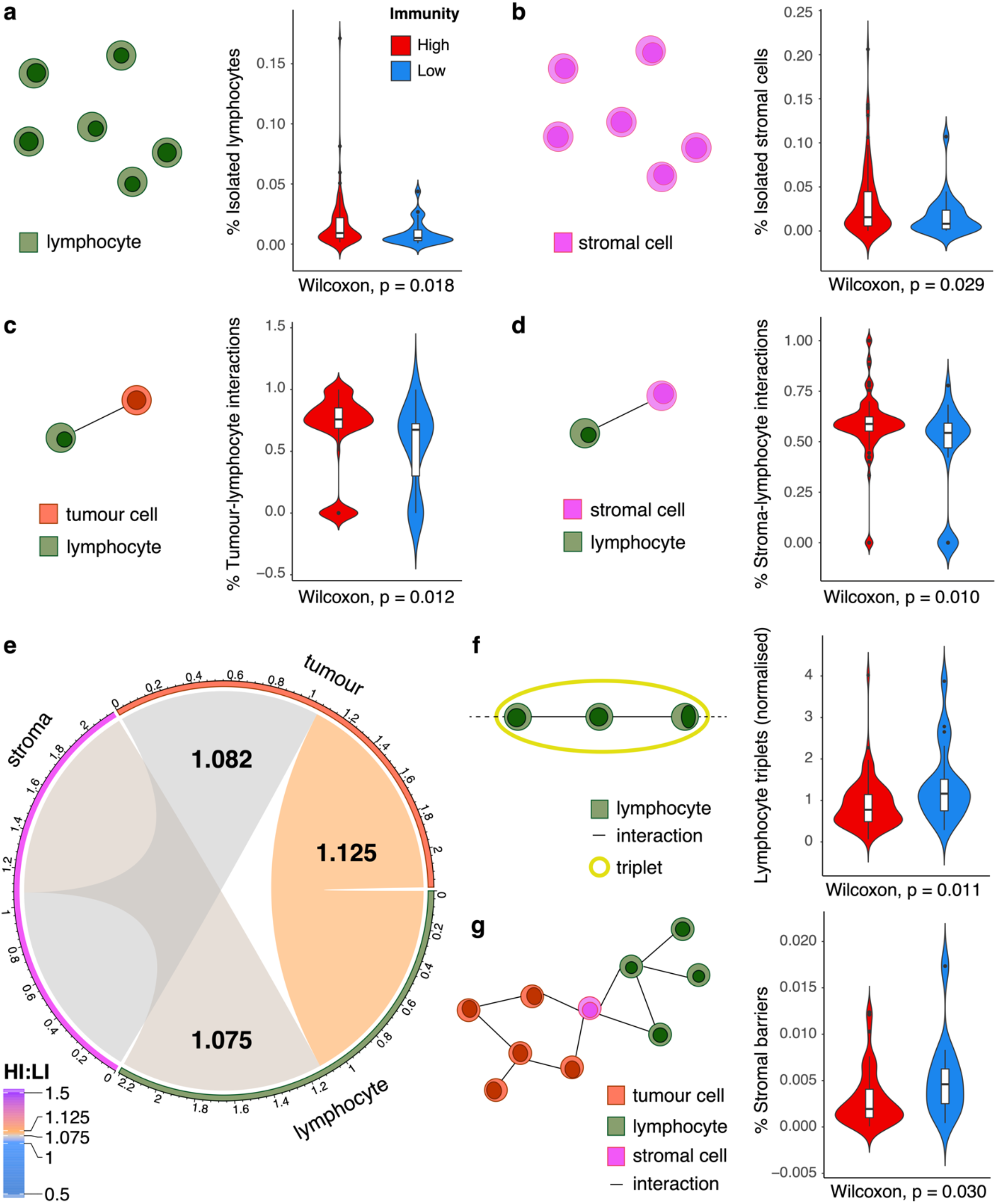
Cellular organization differences in high versus low immunity tumours. High (red) and low (blue) immunity groups are compared in terms of: (a) the fraction of isolated lymphocytes; (b) the fraction of isolated stromal cells; (c) the fraction of direct tumour-lymphocyte interactions; (d) the fraction of direct stroma-lymphocyte interactions. (e) The fold change in interactions established between pairs of cell types in high versus low immunity tumours. The ratio of median numbers in either group is depicted. (f) Fraction of lymphocyte triples compared between high and low immunity samples containing at least one such structure. (g) Fraction of stromal barriers compared between high and low immunity samples containing at least one such structure. Schematic depictions of cellular structures are displayed alongside each comparison.

### Validation of deep learning predictions using spatial transcriptomics

Finally, to validate our model predictions using an orthogonal approach, we employed spatial transcriptomics data available for one colorectal tissue slide from the Visium platform (Figure 6a). We applied our AI model on the H&E stained image to obtain predictions of the overall immunity within the tumour at patch level (Figure 6b). In parallel, we analysed the spatial gene expression profiles across multiple spots within the image and derived an immunity map outlining the distribution of immune ‘hot’ and ‘cold’ islands across the tissue (Figure 6c-d).

**Figure 6.**
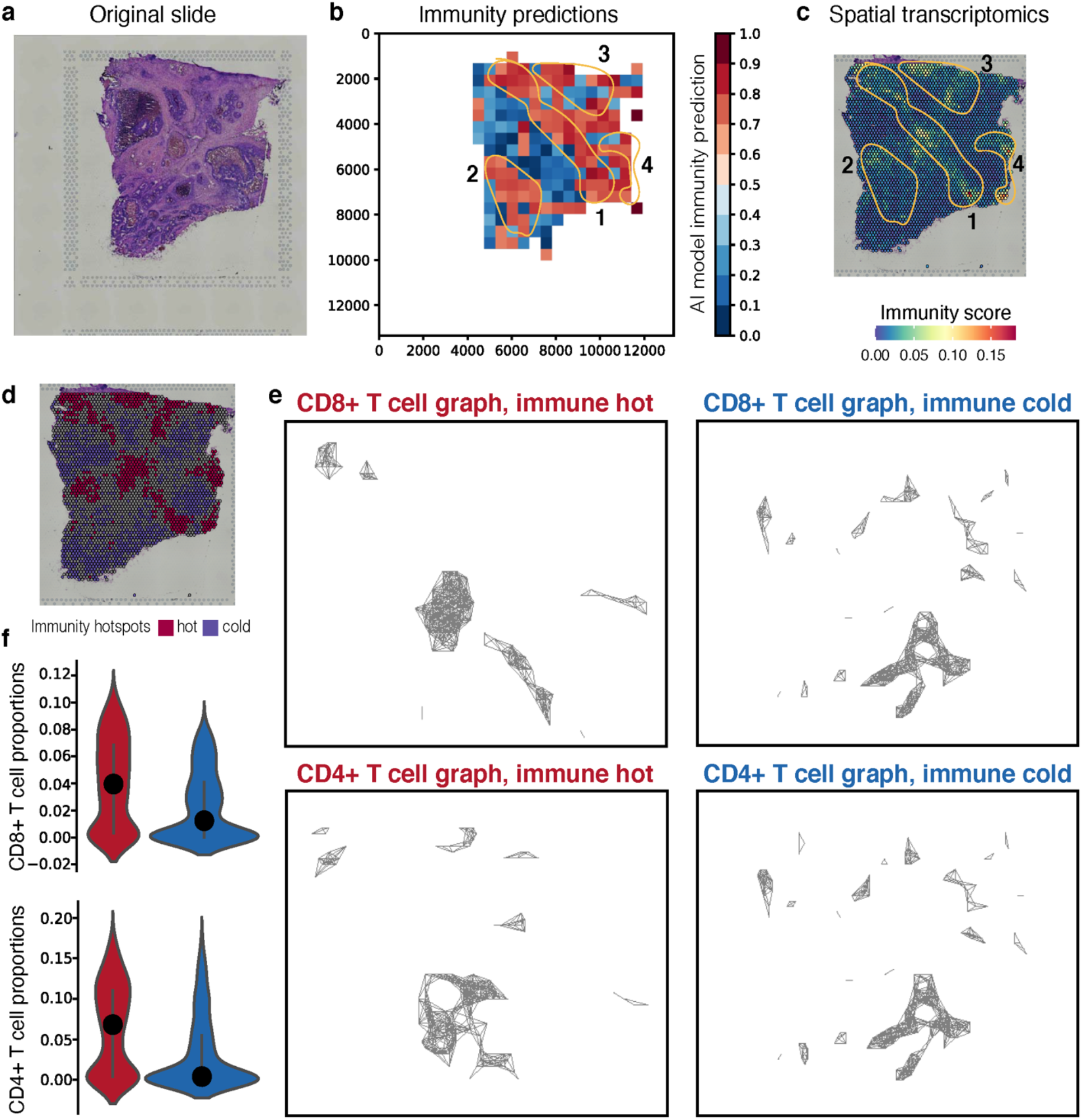
Validation of AI model predictions using spatial transcriptomics data. (a) Original colorectal tissue slide employed for spatial transcriptomics from Visium. (b) The immunity prediction of the deep learning model on H&E staining-derived patches. Red indicates areas of high immunity, blue indicates low immunity. Four high immunity islands are indicated with an orange outline and numbered 1-4. (c) Immunity score across spatial transcriptomics spots. Red and yellow areas indicate high immunity, and the same islands 1-4 as in (c) are indicated on the slide. (d) Immune hot (red) and cold (blue) hotspots defined from spatial transcriptomics. The grey spots have intermediate levels and cannot be classed in either group. (e) CD8+ (top) and CD4+ (bottom) T cell interaction graphs within hot (left) and cold (right) immunity areas. (f) CD8+ (top) and CD4+ (bottom) T cell proportions compared between immune hot (red) and cold (blue) graphs.

By visually comparing the AI model predictions with the spatial profile, we can see there is a good agreement between the two methods (Figure 6b-d). In particular, it is striking that the central area of high immunity (island 1) appears as a diagonal stripe both in the spatial transcriptomics as well as in our model’s predictions, with smaller islands of increased immunity present to the left and right of this region (islands 2-4, Figure 6b-c). The immune ‘hot’ areas presented marked CMS4 features, while the colder areas more frequently contained CMS1/2/3 types of cells (Supplementary Figure 2a-d), as previously shown in the bulk data.

When reconstructing cell-cell interactions within the spatially profiled slide, we confirmed an increase in epithelial-lymphocyte interactions in the immune ‘hot’ compared to the immune ‘cold’ areas (20% versus 5%). Furthermore, interactions between stromal cells and lymphocytes appeared confined to the immune hot regions (41%), with no stromal cells detected in the immune cold areas. The CD8+ and CD4+ T cell spatial structure was very similar in the immune cold regions, with larger interaction modules observed in the immune hot regions (Figure 6e). The immune hot regions were denser, with CD8+ and CD4+ T cell graphs having a density metric of 0.095 and 0.073 respectively, and immune cold regions with a density metric of 0.046 and 0.0451. Immune hot CD8+ and CD4+ graphs also presented increased connectivity (12 connected components each versus 30 and 31 connected components in immune cold regions, respectively). As expected, the high immunity areas also presented increased CD8/CD4+ T cell and stromal cell abundance, while containing fewer NK cells (Figure 6f, Supplementary Figure 2e-f).

All these point towards the co-existence of immune hot and cold areas within tumours that display distinct spatial interactions and confirm many of the features we were able to capture through our deep learning and graph models. While this analysis is limited by the availability of a single slide, it serves as a proof of concept that H&E-based deep learning models could be validated using spatial transcriptomics.

## DISCUSSION

In this work, we have employed state-of-the-art methodology to establish an RNA-seq-derived immune signature in colon adenocarcinoma that is prognostic and links with differential efficacy of various chemotherapeutics. We have shown that this signature is detectable in H&E-stained colon cancer tissue. Furthermore, we have introduced novel methods to explore the ample space of cellular interactions underlying distinct tumour immunity phenotypes, unveiling specific rewiring that could inform diagnosis and treatment.

The encouraging performance of 82% for our tumour immunity classifier in H&E images suggests that integrating such images and transcriptomics data could support faster pathology annotation and triage in a setting where this staining procedure is routine. Our model’s performance was similar to that of other methods assessing immunity-linked phenotypes like MSI, mutability or methylation status in colorectal cancer^25^, and outperformed immune classification/immunotherapy response models in breast cancers^40^ and melanoma^41^ with AUCs of 76-78%. It is worth noting that multiple studies have applied deep learning for lymphocyte feature extraction in colorectal cancer and linked this with outcome^42,43^. However, these fall short of providing a direct H&E-based classifier. While our model does not outperform MSI classifiers developed in this cancer^44,45^, it captures a different and more versatile phenotype of tumour immunity. It also helps us explore the limits of prognostic classification based on expression-derived signatures, for which datasets are much more widely available.

We also showed that not all T cells associated with the antitumour immune response are equally detectable in the cancer tissue when trained on expression markers. This could be due to morphological confounders or to the expression signatures not being specific enough for some cell types. Nevertheless, the cells effectively responsible for cytotoxicity (effector CD8+ and central memory CD4+ T cells) presented the best classification performance, suggesting that both short-term and longer-term immune stimulation may be captured.

Our investigation of the cellular organisation of the tissue has highlighted niche structures such as dense lymphocyte clusters/triplets, or stromal barriers which may account for the lack of immune recognition and worse prognosis in the low immunity group, as reported by other studies too^18,46^. This showcases the importance of spatial analysis of the tumour microenvironment to understand cancer progression. A limitation is that the cell types identified are rather generic. Improved methods are needed in the future to distinguish diverse cell populations, e.g. cancer-associated fibroblasts, T and B cell subsets, and gain a finer-grained resolution of the landscape of cell-cell relations established. Moreover, the structures studied here should be further investigated experimentally to clarify the mechanism by which they contribute to immune evasion. Finally, our spatial transcriptomics validation illustrates one key factor that needs to be built into such models in the future: the spatial heterogeneity of immune hot/cold phenotypes. Future studies should focus on integrating deep learning on H&E slides, spatial transcriptomics models and graph reconstruction methods to obtain a spatially-informed predictor of tumour immunity and response to therapies.

This study proposes a prognostic classifier for tumour immunity in colon adenocarcinoma with distinct tumour and microenvironment architecture features. While we have based our classifier on expression rather than protein-level/IHC data, our immune signature is nevertheless highly prognostic and many of its features are recapitulated in spatial transcriptomics data. It thus could be valuable in the clinic as additional support for treatment decisions. Most importantly, we propose a novel integrative approach to digital pathology analysis in cancer, combining H&E-stained slides and matched RNA-seq data through deep learning, and making use of the capabilities of the Neo4J graph database methodology to efficiently quantify and explore tissue landscapes and cellular interactions. This framework enables a faster, more extensive and more interpretable exploration of key immunity features than with traditional approaches, and could be easily adapted to answer a variety of biological questions in cancer as well as healthy tissue settings.

## MATERIALS AND METHODS

### Molecular data sources and immune stratification

RNA-seq data from 456 colon adenocarcinoma (COAD) tumours, along with clinical and pathology information, were retrieved from The Cancer Genome Atlas (TCGA) using the *TCGAbiolinks* R package. No samples were excluded based on demographics criteria. We estimated the relative abundance of various lymphoid and myeloid cell subsets, endothelial cells and fibroblasts (Supplementary Table 1) based on the expression of cell type-specific markers using ConsensusTME^28^. The ‘immunity score’ was defined as the average abundance across all cell types within a sample, as in the original study (Supplementary Figure 1a). This score was used to stratify the cohort into two groups representing low and high immunity. For this, we used the threshold that maximises the difference in overall survival between the high/low immunity groups, i.e. testing different thresholds by sequentially increasing their value using the *survminer* R package.

xCell^27^ expression-based estimates of CD8+ and CD4+ naïve, central and effector memory T cell populations were obtained for all TCGA COAD cancers from https://xcell.ucsf.edu/. The cohort was split by the mean infiltration estimate of each CD8/CD4+ T cell population.

Signatures of intratumoural NK cell activity were assessed based on the expression of NK cell receptors, tumour ligands and cytokines as detailed in Huntington et al^30^. An expression score summarising these activities was defined per sample using single sample Gene Set Enrichment Analysis via the *GSVA* R package.

We derived the molecular phenotypes of colorectal cancer from Liu et al^31^ and Guinney et al^32^, and retrieved the predicted drug sensitivity IC50 values for TCGA samples from Li et al^47^.

### Image pre-processing

A total of 874 images of H&E-stained tissue collected from 456 COAD patients were obtained from the TCGA Genomic Data Commons Data Portal (GDC Data Portal) (RRID:SCR_014514, https://portal.gdc.cancer.gov/). Because of the high resolution and large scale of these images, a common pre-processing method before applying deep learning approaches for the classification of whole slide images (WSIs) is to crop them into small sections called tiles^64^. We extract all possible non-overlapping tiles following a grid structure and we filter those including more than 50 percent of background. We set the size of the tile to 512px by 512px, yielding a total of more than 2 million of them, which were then used to train and test our model (described below).

To avoid inconsistencies in the preparation of histology slides arising from dye concentration, duration of staining and temperature differences^48^, we employed StainTools (https://pypi.org/project/staintools/) to normalize each H&E patch used in this study. The stain matrix estimation was set to be calculated via the Vahadane method.

### H&E-based classifier of tumour immunity

To classify immunity levels in H&E images, we used a model consisting of two parts: a convolutional neural network (CNN) feature extractor followed by a non-linear classifier (Figure 2a). We based the feature extractor backbone in the InceptionV3 architecture^43^. First, we resized the 512×512px tiles with three colour channels to 299×299px, as this is the required input size of the model. Furthermore, we scaled the input pixel values that were initially in the range of (0;255) to a range of (−1;1).

We removed the top layer of the InceptionV3 original architecture, and used the bottleneck layer’s feature representation. This converts each input image of size 299×299×3 into an 8×8×2,048 block of features. Here, we average over the 8×8 spatial locations, using a Global Average Pooling 2D layer to convert the block of features to a single 2,048-element vector per image. We feed this vector image to a fully connected classifier to convert these features into a single prediction per image. It consists of two dense layers of 1,024 and 512 units, respectively, with a RELU activation function. We applied a dropout regularisation to the output of the first dense layer. Low immunity samples are predicted as class 0 and high immunity as class 1.

We initialised the parameters of the InceptionV3 layers with weights trained in the ImageNet dataset^49^. To avoid destroying the pre-loaded weights, we trained the full model end-to-end with a small learning rate (1e-5). In this way, we fine-tuned the higher-order feature representations in the base model to make them more relevant for this task. During the training, we introduced sample diversity by applying random transformations to the input images, such as rotation, shearing, zooming, horizontal and vertical flipping. To avoid overfitting, we applied L2 regularisation to the kernel of the 2D convolutional layers of the InceptionV3 model during the optimisation. We added this penalty to the loss function as the sum of the squared weights.

We used 70% of the samples for training and 30% for testing, undersampled to the lower class. We repeated each experiment five times with different chosen random samples for training and testing. To present the results, we show the Receiver Operating Characteristic (ROC) curve and the Area Under the Curve (AUC) for each experiment and the total average for all the cross-validation splits.

### Graph-based reconstruction of cell-cell interactions

We used the HoVer-Net computational pathology pipeline trained on the CoNSeP dataset^37^ to segment and classify nuclei within H&E-stained tiles into four categories depending on cell type: tumour cells, lymphocytes, stroma and miscellaneous cells. The miscellaneous category groups artefacts or ambiguous cell types e.g. necrotic, mitotic cells and others that cannot be categorised. This category was discarded from further analysis.

The identified nuclei and their positioning within the tissue were used to reconstruct and analyse the spatial interactions between cells. Each nucleus/cell was represented by a node in a graph. We determined interactions between cells based on spatial proximity, with any two cells situated <35 μm apart assumed to be interacting^50^. We assigned an edge between adjacent cells, depicting the interaction. We then employed the Neo4J Graph Database framework (https://neo4j.com/) to store and efficiently query the graphs derived from the WSIs of 110 patients belonging to either the high or low tumour immunity class.

We compared the cell type abundance and the frequency of different interactions between the high and low immunity groups. A *stromal barrier* was defined as an instance where lymphocytes can reach a tumour cell by crossing a stromal cell in each sample. *Lymphocyte triplets* were defined as three lymphocyte cells sequentially connected. We normalised the number of stromal barriers by the sample’s total number of stromal-stromal relations. Similarly, we normalised lymphocyte triplets by the sample’s total number of lymphocyte-lymphocyte relations.

### Spatial transcriptomics data analysis

The human colorectal cancer patient sample was downloaded from 10x genomics (https://support.10xgenomics.com/spatial-gene-expression/datasets). The output from the Space Ranger Visium pipeline was used for analysis. The *SCTransform* R package was used to normalise the data using a regularised negative binomial regression method. The *Seurat* R package was used to calculate and visualize the gene module scores across the slide. Immunity was scored for each spot using the ConsensusTME methodology. Independently, cell type and state proportions for each spot were estimated using the DestVI package. DestVI requires scRNA from the same tissue for deconvolution. 18,409 cells from 2 colorectal patients were downloaded from Lee et al^51^. The major cell types consisted of B cells, T cells, Epithelial cells, Mast cells, Myeloids and Stromal cells, consistent with the HoVer-Net cellular deconvolution categories. To further break down the cellullar categories, we also used the minor class labels for CD19+CD20+ B, CMS1, CMS2, CMS3, CMS4, IgG+ Plasma, Lymphatic ECs, Myofibroblasts, NK cells, Proliferative ECs, Smooth muscle cells, Stromal 1, Stromal 2, T follicular helper cells, T helper 17 cells, Tip-like ECs, cDC cells as labels. Scanpy^52^ (Single-Cell Analysis in Python) and Squidpy^53^ (Spatial Single Cell Analysis in Python) packages were used for graph analysis. This included graph visualisation and graph metric algorithms. Immune hotspots were calculated from the immune score signature using PySAL (Python Spatial Analysis Library) and separate immune cold and immune hot graphs were calculated from these immune hotspot regions.

### Statistics

Cell organization and disease characteristics were compared between groups using the Wilcoxon rank-sum test. The association between immunity groups and patient outcomes was evaluated using Cox proportional hazards models.

## Data availability

The results published here are based in part upon publicly available data generated by the TCGA Research Network (https://www.cancer.gov/tcga). All these data comply with ethical regulations, with approval and informed consent for collection and sharing already obtained by the TCGA consortium.

The spatial transcriptomics data employed in the study was freely available for reuse from 10x Genomics through the Visium platform (https://support.10xgenomics.com/spatial-gene-expression/datasets).

Ethical approval and written informed consent were not required for this study.

## Code availability

The code developed for the purpose of this study can be found at the following repository: https://github.com/secrierlab/TumourHistologyDL

## AUTHOR CONTRIBUTIONS

MS designed the study and supervised the analyses. MPC built the deep learning classifiers, performed the nuclear segmentation and classification, built the Neo4J graph database and analysed the interactions in HI and LI tumours. GMT and MPC developed the immunity signatures. GMT and MS performed associations with molecular subtypes of colorectal cancer, and further interaction analyses. EW performed the spatial transcriptomics analysis. SP performed the Visium slide TIFF image conversion and pre-processing. All authors wrote and approved the manuscript.

