## Supplementary Material for "A deep learning and graph-based approach to characterise the immunological landscape and spatial architecture of colon cancer tissue"

**Supplementary Table 1. Immune infiltration estimates from bulk RNA-seq data obtained from the TCGA COAD cohort.** The estimates reflect relative enrichment or depletion of a specific cell population within the tumour and have been obtained using ConsensusTME.

**Supplementary Video 1. Example of graph navigation in the Neo4J database of cellular interactions to uncover a stromal barrier.** The video illustrates the interactivity capabilities of Neo4J using cell interaction data that has been derived from cancer H&E slides of colon adenocarcinoma.

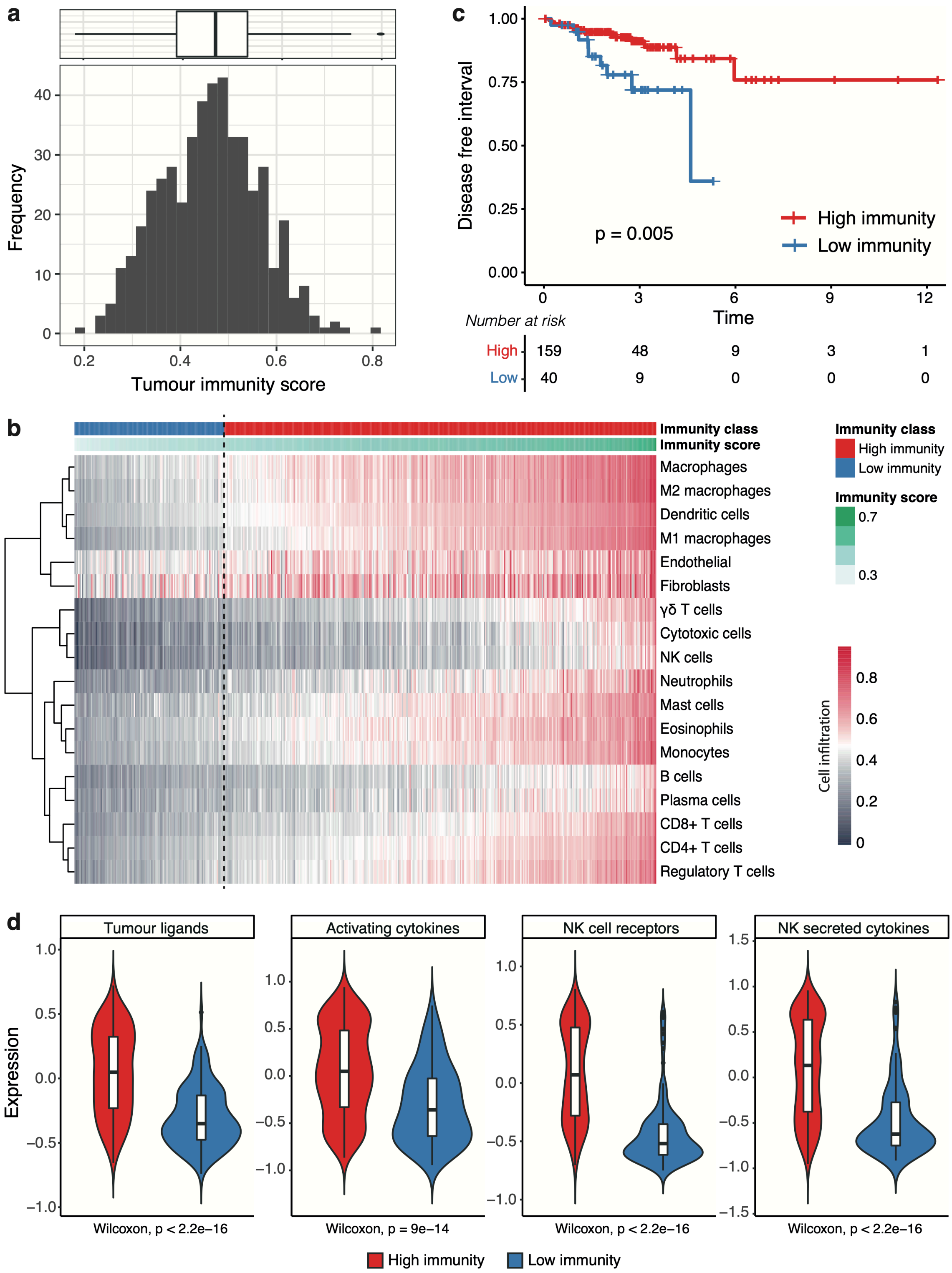


**Supplementary Figure 1. Tumour immunity scores and characteristics.** (a) Distribution of immunity scores calculated using ConsensusTME. (b) Heat map illustrating the abundance of different cell populations within samples across the entire cohort studied. The cell infiltration is estimated using ConsensusTME. The overall immunity score and the binary immune classification (using the 0.39 cut-off) are also indicated. (c) The high and low immunity groups defined using the 0.39 cut-off display significantly different disease free intervals. (d) NK tumour ligands, receptors, activated and secreted cytokines compared between the high and low immunity groups.

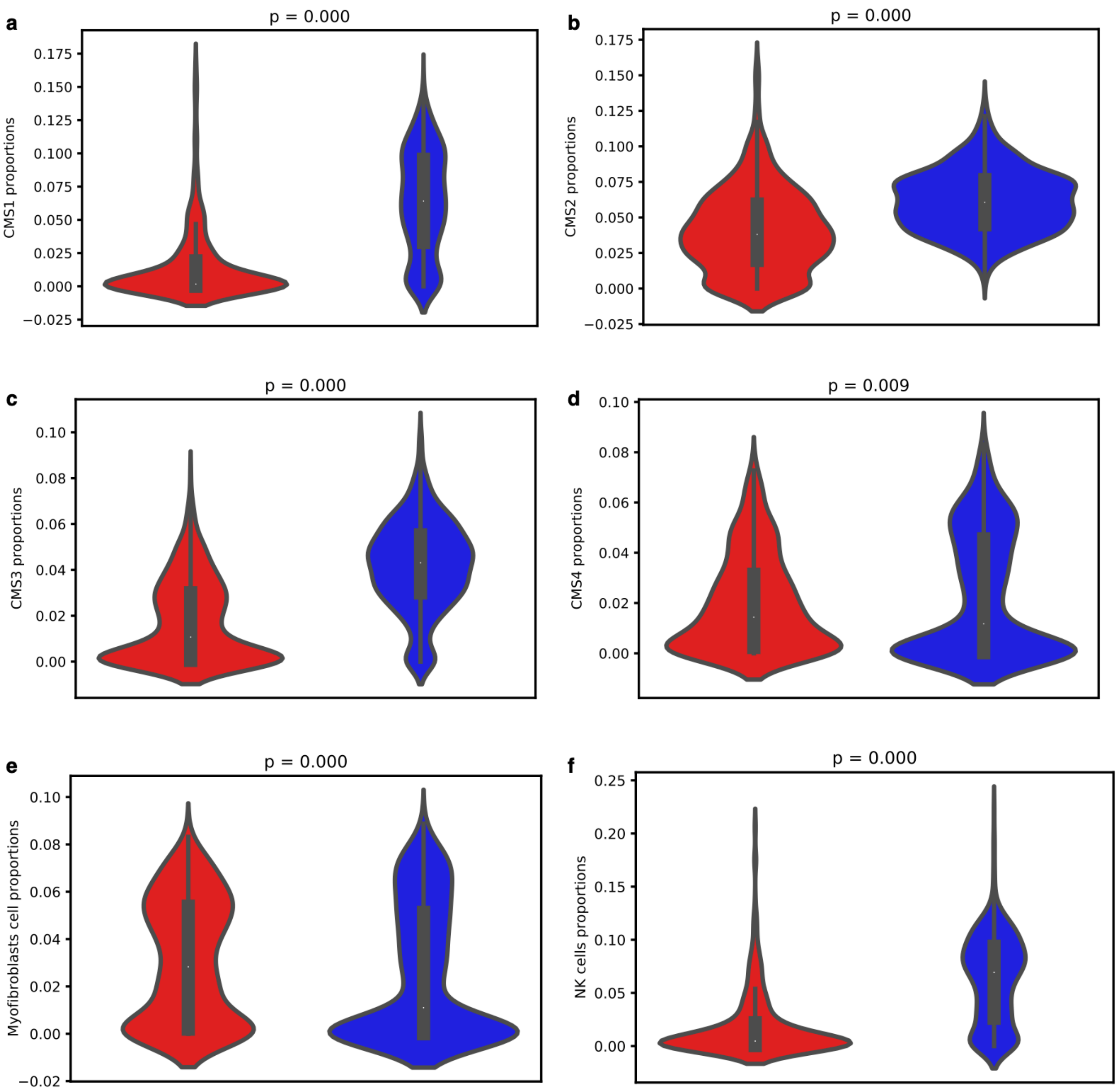


**Supplementary Figure 2. Cell type abundance compared between immune hot and cold regions inferred from spatial transcriptomics data.** The following cell types showed significant differences in abundance between immune hot (red) and cold (blue) regions: (a-d) CMS1-4 tumour cells, (e) myofibroblasts, (f) NK cells.
